# Age-related brain mechanisms underlying short-term recognition of musical sequences: An EEG study

**DOI:** 10.1101/2023.03.12.532256

**Authors:** M. Costa, P. Vuust, M.L. Kringelbach, L. Bonetti

**Author notes:** Corresponding authors: Leonardo Bonetti,; Marco Costa.

## Abstract

Recognition is the ability to correctly identify previously learned information. It is an important part of declarative episodic memory and a vital cognitive function, which declines with ageing. Several studies investigated recognition of visual elements, complex images, spatial patterns, and musical melodies, focusing especially on automatic and long-term recognition. Here, we studied the impact of ageing on the event-related potentials using electroencephalography (EEG) associated with short-term recognition of auditory sequences. To this end, we recruited 54 participants, which were divided into two groups: (i) 29 young adults (20-30 years old), (ii) 25 older adults (60-80 years old). We presented two sequences with an interval of a few seconds. Participants were asked to state how similar the second sequence was with regards to the first one. The neural results indicated a stronger negative, widespread activity associated with the recognition of the same sequence compared to the sequences that were transposed or completely different. This difference was widely distributed across the EEG sensors and involved especially temporo-parietal areas of the scalp. Notably, we reported largely reduced neural responses for the older versus young adults, even when no behavioral differences were observed. In conclusion, our study suggests that the combination of auditory sequences, music, and fast-scale neurophysiology may represent a privileged solution to better understand short-term memory and the cognitive decline associated with ageing.

## Introduction

Recognition is the ability to correctly identify previously learned items and objects. It is an important part of declarative episodic memory and a vital cognitive function ^1^. Recognition memory allows us to distinguish novel from previously learned information, sometimes effortlessly and in a fraction of a second ^2^. Notably, performances in recognition tasks decline with age ^3-7^, suggesting their relevance for studying ageing and its neurobiological correlates.

In the past decades, the neuroscientific community has directed considerable research efforts toward the investigation of the neural substrate of recognition in the auditory domain. For instance, research on processing for auditory sequence and music has largely focused on the fast-scale, automatic brain responses in relation to deviations inserted in coherent sequences of sounds. These studies revealed the fast pre-attentive neural responses which imply the existence of sensory auditory memory, such as N100 and mismatch negativity (MMN) ^8-14^. Remarkably, it has been shown that these brain processes are affected by ageing. For instance, Pekkonen reported reduced MMN in elderly individuals compared to young controls ^5^. A similar result was presented by Cheng and colleagues ^3^ who showed a reduction in the fronto-temporo-parietal activity underlying MMN in elderly compared to young participants.

Other studies focused on conscious recognition processes. For instance, it has been suggested that by recruiting the dorso-lateral prefrontal cortex, the brain actively recognizes sequential regularities underlying a sequence of items ^15, 16^. Relevant contributions in the investigation of the brain correlates of the auditory system also came from speech processing. For instance, it has been shown that the simultaneous role of gamma and theta bands during auditory processing was crucial for the understanding of spoken human languages ^17, 18^. In an additional study on speech recognition, it has been revealed that while theta activity was mainly restricted to the auditory cortex, delta activity originated in downstream auditory regions ^19^. Previous studies also investigated the impact of ageing on the neural mechanisms underlying speech recognition. For instance, Tremblay, Brisson and Deschamps ^6^ found that, even after accounting for hearing thresholds and auditory attention, speech perception declined significantly with age. The decline was associated with thinning of the cortex in regions involved in auditory and speech processing, as well as executive control. Moreover, older adults showed reduced brain responses in the right superior temporal cortex and increased responses to noise in the left anterior temporal cortex. Moving to research on music neuroscience and sound encoding, it has been discovered that the first 220 ms of sound processing recruited a large network of connected brain areas, especially in the right hemisphere, such as Heschl’s and superior temporal gyri, frontal operculum, cingulate gyrus, insula, basal ganglia, and hippocampus ^20^. Remarkably, these brain areas were equally central within the whole brain network, even if auditory cortex and insula were characterized by a much stronger activity than the other areas. Similarly, long-term recognition of short musical sequences recruited nearly the same brain network. However, in this case, the recruitment was bilateral, and it showed hierarchical dynamics from lower- to higher-order brain areas in different frequency bands (e.g. 0.1-1 Hz and 2-8 Hz) ^21-24^. In addition, previous electroencephalography (EEG) research examined the induced responses underlying recognition of auditory stimuli. Authors reported increased gamma power and phase locking between auditory areas when participants successfully recognized familiar stimuli such as well-known environmental sounds ^25, 26^. In the neurobiology of ageing research, several studies suggested that the brain functioning during long-term memory tasks is impaired in older compared to young adults. For instance, Gajewski and Falkenstein ^27^ revealed decreased and delayed ERP components (e.g. N200, P300a and P300b) in older adults when they performed a two-back memory task. Along this line, Vaden, Hutcheson, McCollum, Kentros and Visscher ^7^ showed that in a suppression of visual information task, the correct performances were associated with a robust modulation of alpha power only in young but not in older adults. Similarly, Federmeier, McLennan, De Ochoa and Kutas ^4^ demonstrated that older compared to young adults had a reduced neural efficiency when recognizing familiar words.

In addition to long-term recognition, the perception, retention, and recognition of information can also occur for short periods of time. This is the case of short-term memory, which is the ability to retain temporarily limited amount of information in a very accessible state ^28^. It is slightly different from working memory (WM) which consists not only of holding but also of manipulating the online information in an abstract manner ^29^. Both memory subsystems have been mostly investigated in the verbal and visuospatial domain, while less research has been conducted with auditory and musical stimuli. In this context, while a phonological loop was included in the influential memory model of Baddeley and Hitch ^30^, more recent research has proposed a distinct tonal loop for musical information ^31-33^. Daligault and colleagues ^34^ have shown a double dissociation between verbal memory and tonal memory. Further, cases of congenital amusia have revealed impaired memory for tone and timbre sequences but preserved short-term memory for spoken syllable sequences ^35-38^. In a classic functional magnetic resonance imaging (fMRI) study on short-term memory for auditory stimuli, Gaab, Gaser, Zaehle, Jancke and Schlaug ^39^ measured the brain activity of participants while performing comparisons of different simple melodies. When participants successfully performed the task, their brain was mainly active in the superior temporal, superior parietal, posterior dorsolateral frontal, and dorsolateral cerebellar regions, and left inferior frontal gyri. More recently, Kumar, Joseph, Gander, Barascud, Halpern and Griffiths ^40^ used fMRI to show that activity and connectivity between primary auditory cortex, inferior frontal gyrus and hippocampus were necessary for performing an auditory WM task, which consisted of maintaining short sequences of single sounds. WM has also been studied using EEG. For instance, one component that has been repeatedly observed in relation to WM is the negative slow wave (NSW), which is a broadly distributed sustained negativity that persists during the maintenance time in WM tasks ^41, 42^. Ruchkin, Johnson Jr, Grafman, Canoune and Ritter ^43^ showed that this component increased as the memory load increased and was shown reliable for both visuospatial and verbal tasks. Further, Rösler, Heil and Röder ^44^ found that in trials where a larger NSW amplitude was observed during the retention time, there was a higher probability of successfully remembering the information at test, suggesting that the NSW is important for the memory performance on the task. Recent research on short-term memory has also employed magnetoencephalography (MEG). For instance, using musical stimuli, Albouy and colleagues ^35, 45^ investigated the brain activity underlying memory retention. They showed that theta oscillations in the dorsal stream of the participants’ brain could anticipate their skills when performing an auditory WM task using musical stimuli. Interestingly, previous research has also investigated the impact of ageing on short-term recognition of information, showing impaired brain functioning in older compared to young adults. Maylor, Vousden and Brown ^46^ asked participants to immediately recall a list of letters that was previously presented. Authors found that older adults committed a higher number of errors and omission. In a similar study, Gilchrist, Cowan and Naveh-Benjamin ^47^ reported a reduction in the short-term elaboration of information and working memory abilities of older compared to young adults.

Altogether, several studies have been performed on memory recognition. In addition, there are evidence pointing at a decrease in the efficiency of the brain and cognitive abilities of older adults. However, the brain mechanisms underlying processing and short-term recognition of musical sequences are not fully understood yet. In particular, little is known on the differential brain activity underlying recognition of musical sequences that were presented a few seconds earlier versus novel sequences. In addition, the specific impact on ageing on such brain functioning is not known. This is partially due to the fact that while a large number of studies has focused on the biological correlates of Alzheimer’s and other types of dementia ^40, 48, 49^, the investigation of the neural changes associated with normal, non-pathological ageing has received less attention. In addition, among the previous studies which explored the neural correlates of non-pathological ageing, a large focus has been directed toward resting state ^50, 51^ and automatic memory ^3, 5^. Conversely, less research has explored the fast-scale neurophysiological correlates of conscious memory processes such as short- and long-term recognition of auditory sequences. For these reasons, in our study we filled this gap by investigating the electrophysiological signature of short-term recognition of musical sequences in two groups of young and older healthy adults.

## Methods

### Participants

We recruited 54 participants, which belonged to two age groups: young and older adults. The age range of the young participants was 20-30 years old. They were 29 (11 males and 18 females) with a mean age of 23.58 (*SD* = 3.43). Mean age for males was 25.81 years (*SD* = 4.33), and mean age for females was 22.22 (*SD* = 1.80). The age range for the older adults was 60-80 years old with a mean age of 65.24 (*SD* = 5.88). They were 25 (ten males and 15 females). Mean age for males was 64.30 years (*SD* = 6.94), and mean age for females was

65.87 (*SD* = 5.22). Participation was on a voluntary basis. None of the participants had a formal music education and/or music practice experience that exceeded 10 years. None had hearing disabilities or pathologies (self-reported). Twenty-nine (93.54%) were right-handed and two (6.45%) were left-handed.

Every participant was informed about the details of the experimental procedure prior to engaging in the study, and a formal informed consent had to be signed. The study was approved by the Ethics Committee of the University of Bologna (Ref. 27281, 05.02.2021) and was conducted complying with the Declaration of Helsinki – Ethical Principles for Medical Research.

### Procedure

We used a same/different task which presented participants with pairs of musical sequences (S1 and S2). For each pair, participants were asked to state whether S2 was the same or different compared to S1.

First, participants received instructions about the task and signed the informed consent. Then, they were acquainted with the experimental room and the EEG preparation and recording. The participant seated in front of a 19’’ monitor at a distance (eyes-monitor) of about 60 cm, and in front of a loudspeaker at a distance (eyes-loudspeaker) of about 160 cm. A preliminary training session (eight trials) familiarized the participant with the procedure and the stimuli. The experimental task consisted of 240 trials divided in six blocks of 40 trials each (see next section for detailed information). Each block included 10 trials for each of the four conditions. Each trial started with a counter lasting 0.5 s presenting the number of block and the number of trials. The counter was followed by a white fixation cross at the center of the black screen. After one second the first five-tone sequence (S1) was played for a duration of 1.750 s (five tones of 350 ms each). S1 was followed by a silence of two seconds. Then, the second sequence (S2) (duration of 1.750 s) was presented. After the end of the second sequence, the fixation cross disappeared, and a new message invited the participant to rate the degree of dissimilarity between the first (S1) and the second (S2) sequence on a five-point Likert scale ranging from one (completely similar) to five (completely different). Participants rated the similarity by pressing the corresponding key on a keyboard. The maximum time allowed for the response was five seconds. After the response (or after the response was not emitted within five seconds), the screen was left completely empty for an inter-trial interval ranging from 0.5 to 2.5 s. Participants were warmly encouraged not to move and not to move their eyes during the presentation of S1 and S2. The total duration of the EEG recording was about 45 minutes per participant. The total duration of the experimental session (including preparation) was about 90 minutes.

### Stimuli

Stimuli consisted of two sequences (S1 and S2) of five-tone melodies, separated by an interval of 2 s. Each tone had a duration of 350 ms, and each sequence had a duration of 1,750 ms. Tones were generated by a Roland SC-880 sound module, using a flute timbre, with 20 ms rise and decay time. The flute timbre was chosen having a rather flat loudness envelope. Sounds could range from B3 to C#5, for a total of 15 sounds. Their loudness was equalized to 43.4 sons using the Genesis Loudness Toolbox for Matlab (Genesis, 2009), applying the ANSI S34 2007 norm (American National Standards Institute, 2007), and were presented to participants with an average loudness of 68 dB(A), after assessing that the participants could comfortably hear the sounds.

The sequence S2 with regards to S1 in each trial could belong to one of the following four conditions: (a) Same: S2 was completely equal S1; (b) Different: S2 was completely different from S1; (c) Transposed: S2 was transposed one semitone up (50%) or one semitone down (50%) with regards to S1; (d) Last note changed: only the last note of S2 was changed, one-semitone up (50%) or one-semitone down (50%) in comparison to the last note of S1. Each condition included 60 trials, for a total of 240 trials.

To be noted, the repetition of the same note in two following tones was always avoided in order to emphasize a succession of different tones.

### Data acquisition

EEG data was acquired through a Biosemi 64-channel equipment (Biosemi ActiveTwo). In addition to the 64 EEG channels, two sensors were applied to the left and right mastoids, respectively. An additional sensor was positioned below the left eye. In a bipolar montage with the Fp1 EEG sensor, this sensor allowed the monitoring of vertical eye movements and blinks. Horizontal eye movements were instead recorded with the bipolar montage of two sensors positioned in the outer canthi of the left and right eye. The sampling rate was 1000 Hz. Epochs were created for each first and second sequence (S1 and S2) using a time window of -100, 1750 ms. The trigger in correspondence with the first sound presentation for each sequence was set as time 0.

### Data analysis

Behavioral responses were analyzed for dissimilarity ratings, reaction times, and accuracy. These dependent variables were analyzed with a repeated-measure ANOVA design. Greenhouse-Geisser correction was applied to reduce sphericity problems. Independent variables were Age-Group (20-30; 60-80), Condition (four levels: same, different, transposed, last note), and Musicality (two levels: musical and random sequence). Accuracy was assessed with these indexes: (a) percentage of response one in the Same condition; (b) percentage of response five in the Different condition.

EEG preprocessing, and statistical analysis were performed using EEGLAB ^52^, ERPLAB Toolbox ^53^ and Mass Univariate ERP Toolbox ^54^. Data preprocessing followed a standard pipeline, which included rereferencing to linked mastoids, low-pass filtering (100 Hz), high-pass filtering (0.05 Hz), downsampling to 300 Hz. To remove the artifacts caused by eye blinks, muscle, eye movements, and heartbeat activity independent component analysis was used. Contaminated components were first detected by visual inspection and then removed by the data ^55^. The data was then epoched from -100 to 1750 ms with reference to the onset of the first sound of the sequences. The baseline correction was performed by subtracting the averaged pre-stimulus interval (from -100 to 0 ms) from the whole epoch. Artifacts due to massive interference from sources outside the scalp were detecting by using a moving window peak-to-peak algorithm. This excluded all trials in which a peak-to-peak excursion greater than 150 μV was detected in an EEG channel. Moving window width was set to 200 ms, and window step was set to 100 ms.

Then, inferential statistics was computed to address two main research goals: assessing whether the ERPs of S2 for the four experimental conditions were different (i) and whether the age group of the participants modulated the amplitude of the ERPs of S2 (ii).

First, to test the differences between the ERPs to the experimental conditions, repeated measures, two-tailed permutation test based on the tmax statistic ^56^ were computed, using a family-wise alpha level of 0.05. All time points between 0 and 1750 ms and all 64 scalp electrodes were inserted in the analysis. Repeated measures t-tests were performed for each comparison of the experimental conditions (i.e. Same vs Different (i); Same vs Transposed (ii); Same vs Last note changed (iii)) using the original data and 2500 random within-participants permutations of the data.

Second, to detect reliable differences between the ERPs to the second sequence S2 from young and older adults, these ERPs were submitted to an independent sample, two-tailed permutation test based on the tmax statistic ^56, 57^, using a family-wise alpha level of 0.05. Also in this case, all time points between 0 and 1750 ms at all the 64 scalp electrodes were included in the test. 2500 random between-participant permutations of the data were used to estimate the distribution of the null hypothesis (i.e., no differences between the groups). For each permutation, participants were randomly assigned to one of the two groups without replacement.

The permutation tests were chosen for these analyses because they provide much better spatial and temporal resolution than conventional analysis of variance (ANOVA) while maintaining a desired family-wise alpha level (i.e., it corrects for the large number of comparisons). Moreover, the tmax statistic was chosen for this permutation test because it has been shown to have relatively good power for data (like ERPs) whose dimensions are highly correlated ^57^. 2500 permutations were used to estimate the distribution of the null hypothesis as it is over twice the recommended number ^56, 57^ for a family-wise alpha level of 0.05.

## Results

### Behavioral results

#### Dissimilarity ratings

The repeated-measures ANOVA showed a significant effect for Condition: *F*(2.35, 122.37) = 549.14, *p* < .001, η^2^ = .92. Planned comparisons showed that all the four conditions differed significantly from each other. Mean dissimilarity values were 1.31 (*SD* = 0.31) for the Same condition, 2.33 (*SD* = 0.32) for the Last note condition, 3.10 (*SD* = 0.6) for the Transposed condition, and 4.24 (SD = 0.29) for the Different condition (**Figure 1**).

**Figure 1.**
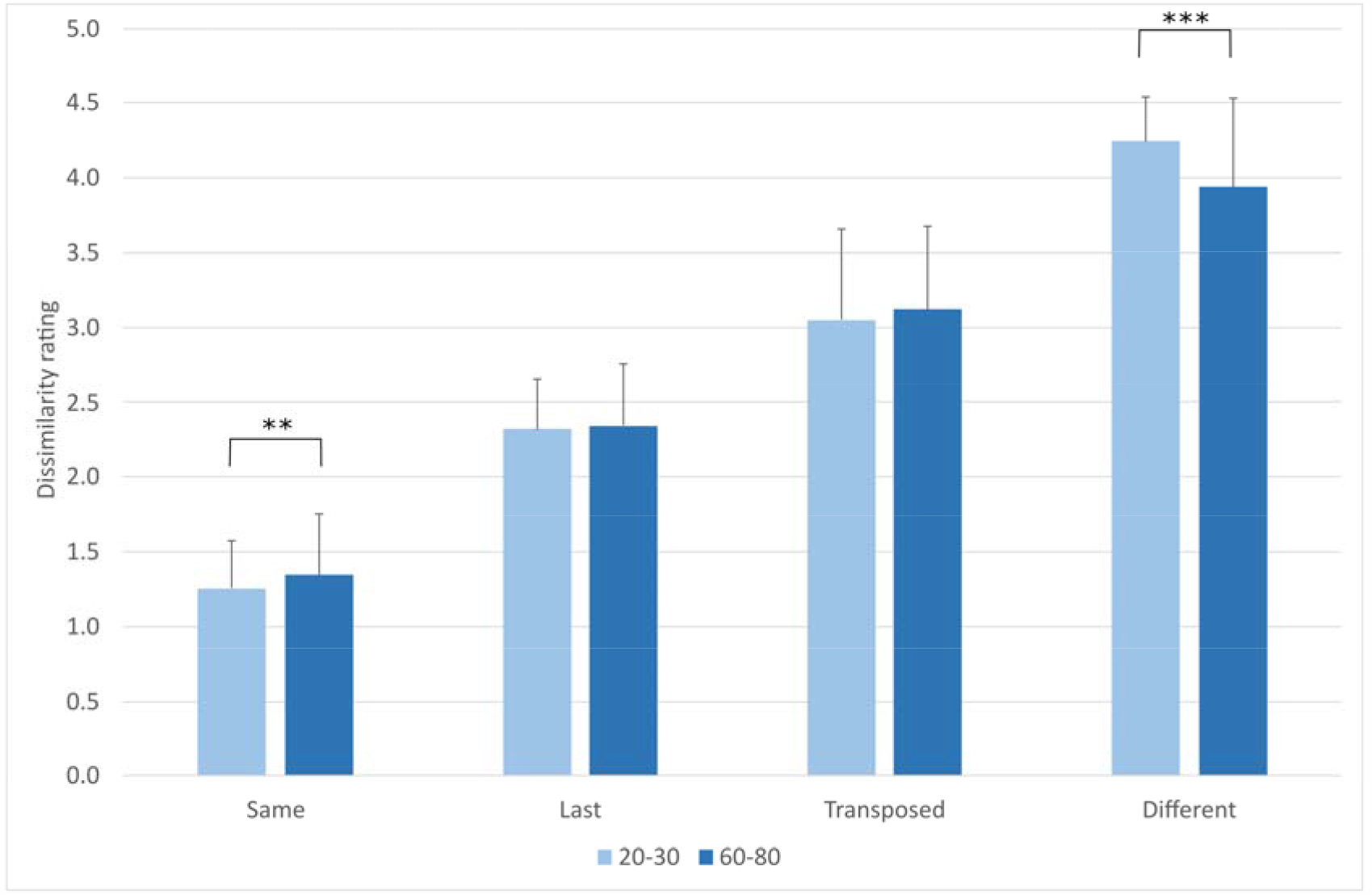
Dissimilarity mean ratings (and standard deviations) for the four experimental conditions.

No significant main effect was found for Age-Group (*p* = .60), but the interaction between Age-Group and Condition was significant: *F*(2.35, 122.37) = 4.45, *p* = .01, η^2^ = .08 (**Figure 1**). Paired comparisons showed a significant difference between the two age groups in the Different condition (*p* = .01) and in the Same condition (*p* = .01). Mean dissimilarity rating for the Different condition was 4.24 (*SD* = 0.29) for young adults and 3.93 (*SD* = 0.59) for older adults. Musicality main effect was not significant (*p* = .21). The interaction between Condition and Musicality was significant: *F*(2.74, 149.77) = 9.94, *p* < .001, η^2^ = .15. The interaction is shown in **Figure 1**, and planned comparisons showed a significant difference for the Same condition (*p* = .008) and the Different condition (*p* = .001). Sex main effect and the interaction with Condition were not significant: *p* = .16 and *p* = .58, respectively.

#### Accuracy

Accuracy related to response one to sequences in the same condition (participants evaluating equal sequences as completely equal) was on average 78.86% (*SD* = 23.26%). Accuracy for the Same sequences was significantly higher in the musical sequences (*M* = 81.20% *SD* = 23.80) than in the random sequences (*M* = 76.51% *SD* = 23.64): *F*(1, 52) = 10.56, *p* = .002, η^2^ = .17. Accuracy for the Same condition was not significantly affected by Age-Group (*p* = .37) and by Sex (*p* = .26). Accuracy related to response five to sequences in the different condition (participants evaluating different sequences as completely different) was 43.82% (*SD* = 21.75) on average. Musicality was a significant factor: *F*(1, 51) = 10.25, *p* = .002, η^2^ = .17. Mean accuracy in musical sequences was *M* = 46.81% (*SD* = 22.04), whereas in random sequences mean accuracy was *M* = 40.84% (*SD* = 21.46). Both Age and Sex were not significant. The accuracy in the Same condition (M = 79.17%, SD = 23.8) was significantly higher than in the Different condition (M = 43.82% SD = 21.75): *F*(1, 51) = 89.29, *p* < .001, η^2^ = .63.

#### Reaction times

The repeated-measure ANOVA showed a significant effect for Condition: *F*(2.64, 137.50) = 46.59, *p* < .001, η^2^ = .47. Mean reaction times were fastest for the Same condition (*M* = 959.79 ms, *SD* = 383.64), and progressively slower for the Last note changed condition (*M* = 1254.77 ms, *SD* = 467.31), the Different condition (*M* = 1318.34 ms, *SD* = 481.65), and the Transposed condition (*M* = 1330 ms, SD = 481.65). Planned comparisons showed significant differences between Same and Last note changed (*p* < .001), between Last note changed and Transposed (*p* = .02), whereas the difference between Different and Transposed conditions was not significant (*p* = .66). The main effect of Musicality was not significant (*p* = .28). The interaction Musicality x Condition was not significant (*p* = .20). Age group main effect was not significant (*p* = .29), as well as sex main effect (*p* = .27), but the interaction between Condition and Age was significant: *F*(2.64, 137.50) = 3.74, *p* = .01, η^2^ = .07. Pairwise comparisons showed that the significance was explained by the difference in the Different condition where the 60-80 group had a mean reaction time of 1183.28 ms (*SD* = 510.23) in comparison to the 20-30 group who had a mean reaction time of 1427.27 ms (*SD* = 356.25). Mean reaction times and standard deviations as a function of the four conditions and age are shown in **Figure 2**.

**Figure 2.**
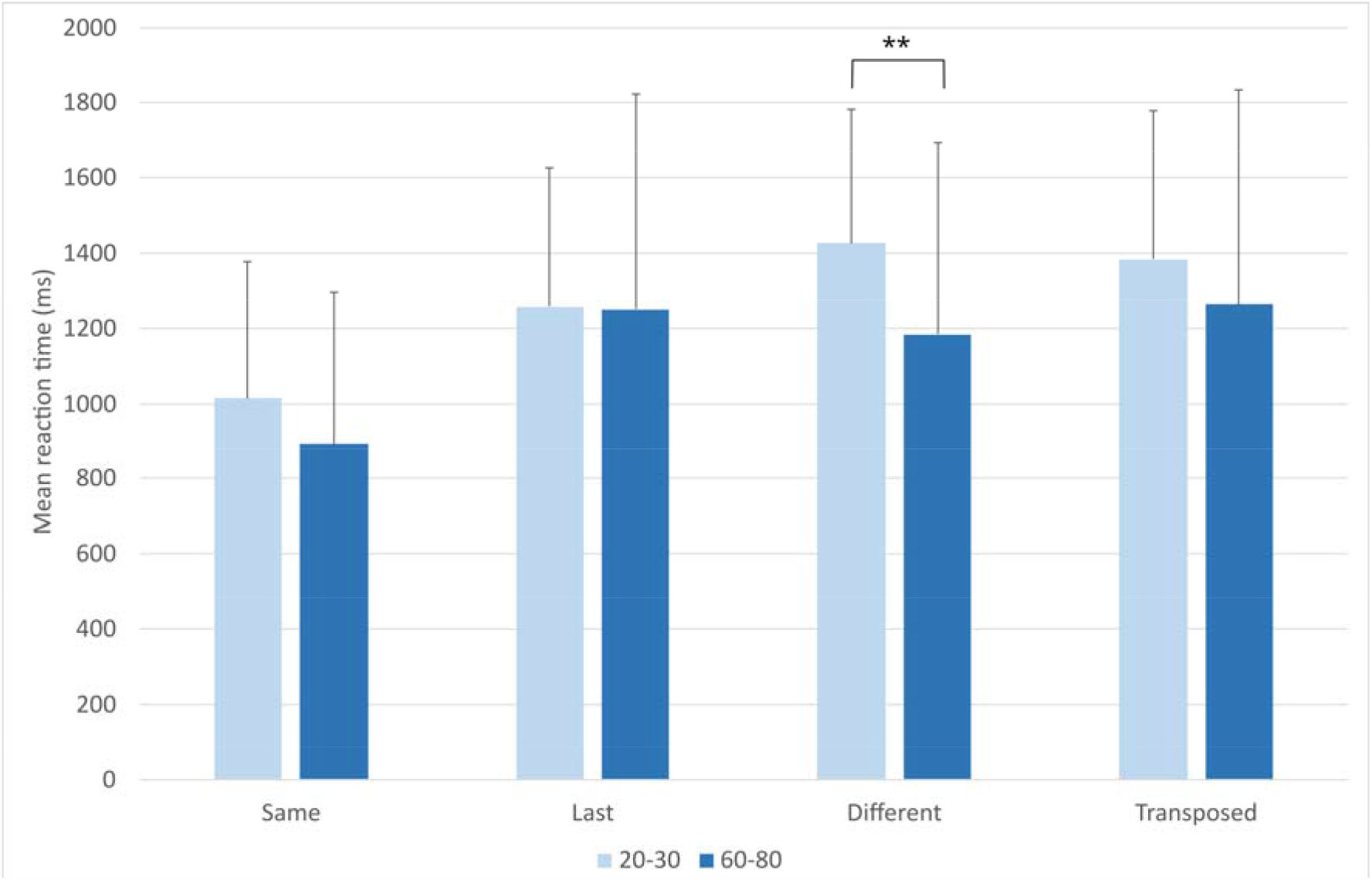
Mean reaction times (and standard deviation) for the four experimental conditions in young and older adults.

### EEG results

#### Same versus Different sequences

Mass univariate analyses in the time window 0-1750 ms with tmax permutation test including Condition (Same versus Different) and Age-Group (20-30 versus 60-80) as independent variables showed significant differences (*p* < .05) between the Same and Different S2 sequences in the following anterior-central sensors: AFz, F1, Fz, F2, F4, FC1, FCz, FC2, C1, Cz, CPz, and Pz), as shown in **Figure 3B**. These channels were inserted into the computation of grand-averaged potentials contrasting Same-Different conditions in young and older adults, as shown in **Figure 3A**. Mass univariate analysis with tmax permutation test including all participants, young adults, older adults and the two age groups are graphically illustrated in **Figure 3C**. The Same-Different contrast was largely significant in young adults, but only in a few time-points with regards to older adults. As shown in **Figure 3A** and **3C**, the main effect of Age-Group was particularly enhanced in the time interval 300-900 ms, especially on the fronto-central sensors Fz and FCz.

**Figure 3.**
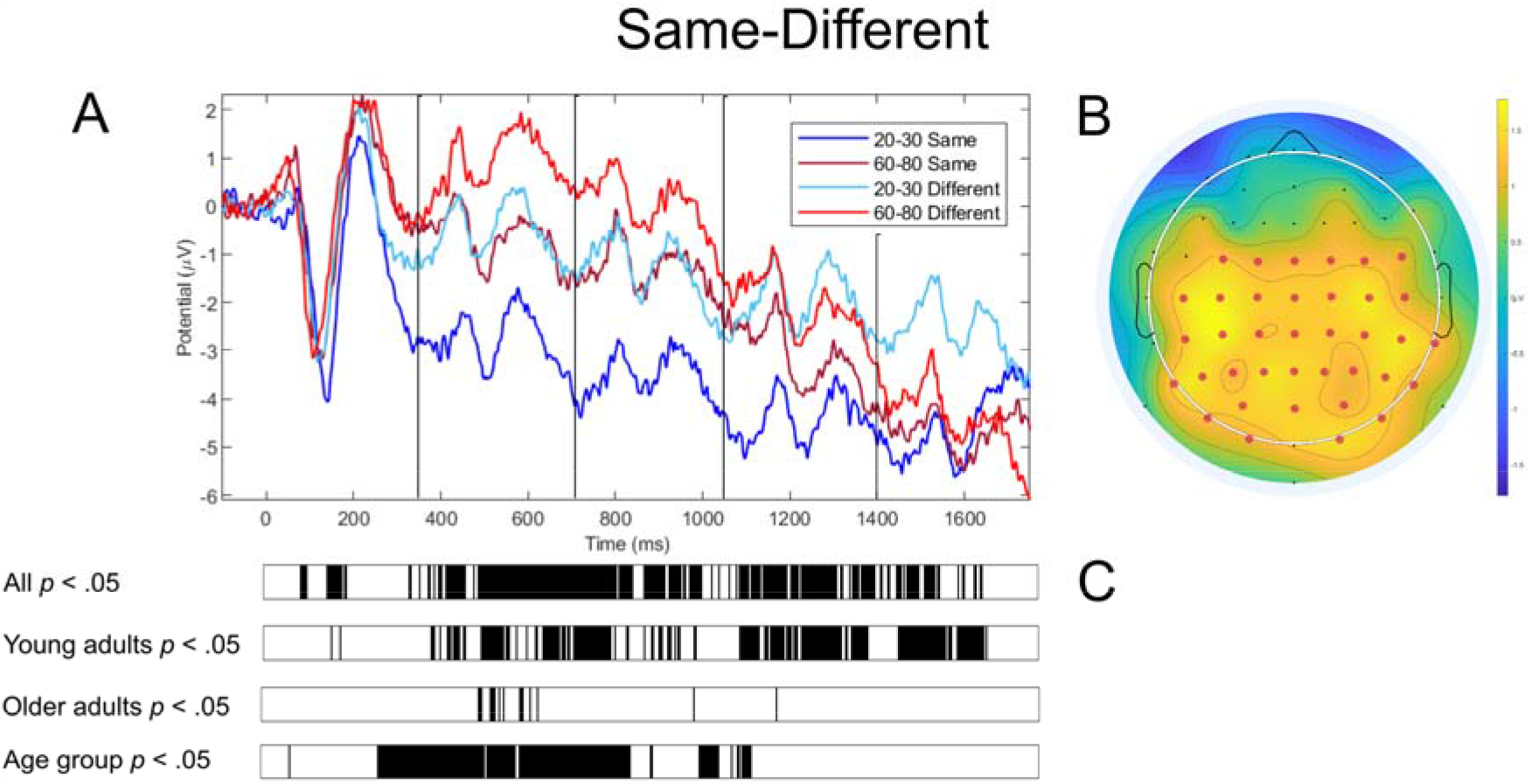
(A) Grand-averaged evoked potential for the conditions Same and Different in young and older adults. (B) Topoplot charting the channels with significant differences in the time interval 0-1750 ms. (C) Bars plotting the significant time bins considering the whole sample, young adults, older adults, and the two age groups. Black lines showed a p < .05.

#### Same versus transposed sequences

Mass univariate analyses in the time window 0-1750 ms with tmax permutation test that included Condition (Same versus Transposed) and Age-Group (20-30 versus 60-80) as independent variables showed significant differences (*p* < .05) between the Same and Transposed S2 sequences in the following central-posterior sensors: C5, CP3, CP1, CPz, CP2, CP6, P7, P5, P3, P1, Pz, P2, P6, PO7, PO3, Poz, PO8, O1, O2), as shown in **Figure 4B**. These channels entered the computation of grand-averaged potentials contrasting Same-Transposed conditions in young and older adults, as shown in **Figure 4A**. Mass univariate analysis with tmax permutation test results regarding all participants, young adults, older adults and the two age groups are graphically illustrated in **Figure 4C**. As shown the Same-Transposed contrast was exclusively significant in young adults but not in older adults. The difference between the two age groups was significant in the time intervals 117-1125 ms.

**Figure 4.**
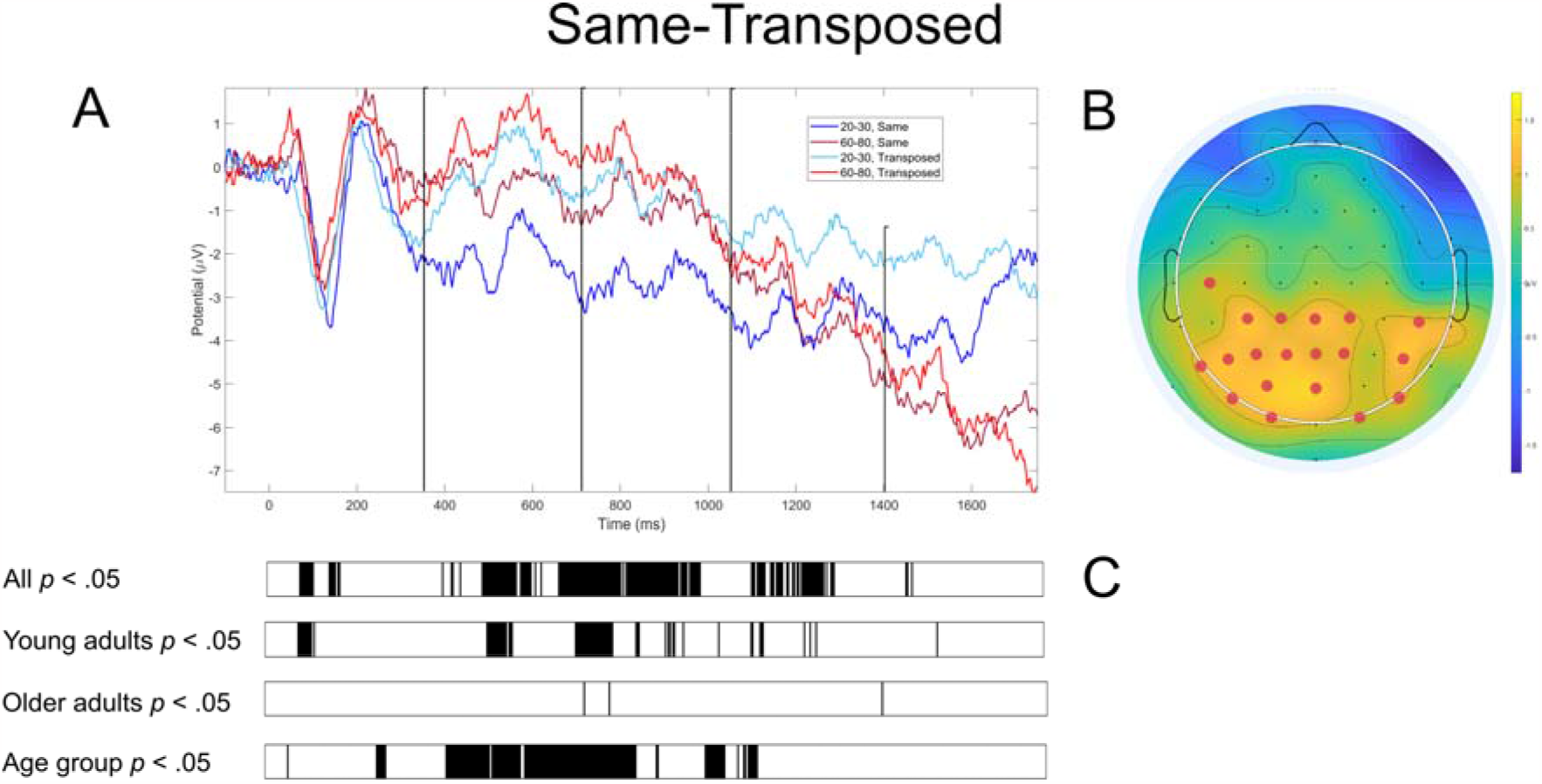
(A) Grand-averaged evoked potential for the conditions Same and Transposed in young and older adults. (B) Topoplot charting the channels with significant differences in the time interval 0-1750 ms. (C) Bars plotting the significant time bins considering the whole sample, young adults, older adults, and the two age groups. Black lines showed a p < .05.

#### Same versus last note

Mass univariate analyses in the time window 0-1750 ms with tmax permutation test including Condition (Same versus Last note) and Age Group (20-30 versus 60-80) as independent variables showed no significant differences in any sensor and any time point. The grand-averaged potentials for the two conditions and distinguishing between young and older adults is reported in **Figure 5**.

**Figure 5.**
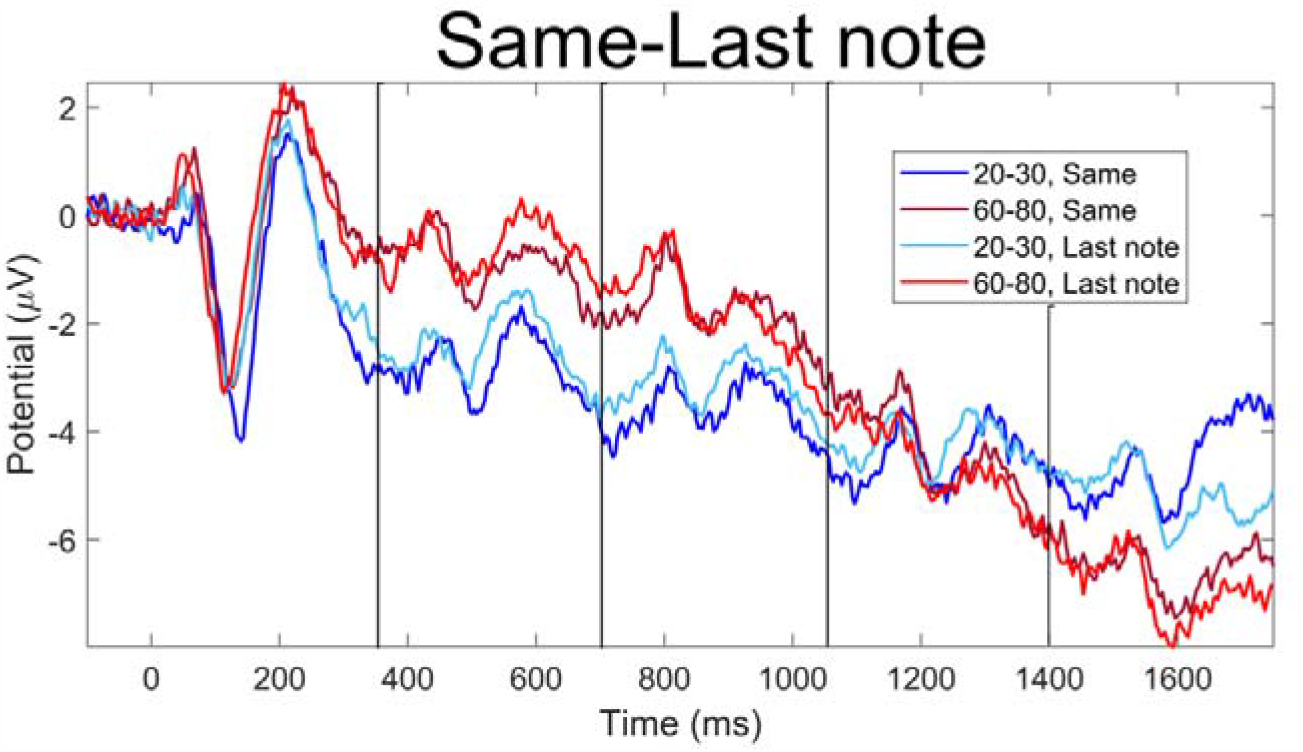
Grand-averaged evoked potential for the conditions Same and Last note changed in young and older adults.

## Discussion

This study showed the event-related potential associated with short-term recognition of auditory sequences in young and older healthy adults. Our paradigm consisted of presenting two sequences with an interval of a few seconds. Participants were asked to state how similar the second sequence was with regards to the first one. Behavioral results showed that similarity ratings were significantly higher for the recognition of the same sequence compared to the sequences that were transposed, completely different, or different for the last tone only. Reaction times were overall faster for the recognition of the same sequence. Moreover, when taking ageing into account, older adults reported overall worse performances than young participants when they had to evaluate the completely similar or different sequences.

The neural results indicated a stronger negative, widespread activity associated with the recognition of the same versus all other categories of sequences. This difference was widely distributed across the EEG sensors and involved especially temporo-parietal areas of the scalp. Notably, the transposed condition was associated to a brain activity in between the same and the completely different sequences. With regards to the impact of age on the brain activity, we reported overall reduced neural responses for the older versus young adults. This was particularly true for the recognition of the same sequences.

The behavioral results are in line with previous studies which reported better scores in recognition tasks for stimuli of lower complexity and degree of dissimilarity ^58-60^. Interestingly, the recognition of the same sequences was associated to faster reaction times, suggesting that the human mind processes familiar items sooner than novel elements. This is also coherent with previous research which highlighted that environmental familiar scenes were more easily processed and recognized than novel ones ^61^, as well as familiar versus novel faces ^62^.

When looking at the impact of ageing on the short-term recognition of sequences, our results showed that older compared to young adults reported worse performances, which is coherent with previous studies. For instance, in a review, Connor ^63^ showed that older adults presented a significantly decreased performance in short-term memory, although she highlighted that the size of such decrement was not particularly large. This is very much in line with the results reported in our study. Differently, Connor reported a higher degree of impaired performance for more complex memory systems such as long-term encoding and recognition. Likewise, Maylor, Vousden and Brown ^46^, asked young and older adults to perform a short-term memory task consisting of immediately recalling a list of letters that was previously presented. They showed that older adults committed a higher number of errors and omission. Similarly, Gilchrist, Cowan and Naveh-Benjamin ^47^ reported a reduction in the working memory abilities of older compared to young adults.

The neural results pointed to an increased negativity associated with the recognition of the previously heard auditory sequences. Although the current study investigated auditory short-term memory, this result is very coherent with previous research ^21-24^ that reported a sustained negativity component associated with the long-term recognition of previously learned musical sequences. In their studies, authors revealed a simultaneous brain processing of the sounds, highlighting fast oscillatory-like responses to each sound (i.e. P50, N100 and P200) and a slower negativity which was particularly marked during recognition of previously learnt music. Additionally, they revealed slightly different dynamical neural networks associated with recognition of complex and simple musical sequences. Along this line, Albouy and colleagues ^35, 45^ showed similar results, mainly concerning WM and memory retention. Indeed, they described a slow negativity associated to an auditory WM task. Moreover, they showed that theta oscillations in the brain regions forming the dorsal stream predicted the participants’ abilities to perform an auditory WM task which required them to maintain and manipulate sound information. In light of these results, we argue that a sustained negativity may be a general neural signature of auditory memory since it has been reported by independent studies for different memory systems (i.e. short-term, long-term recognition, and WM). Future studies are called to investigate short-term recognition with MEG and perform an accurate source reconstruction. In this way, it could be revealed whether the same brain networks are implicated in all these memory processes or if short-term memory has a particular set of brain areas which are differently recruited.

Of particular interest it is the neural response to the transposed condition. In fact, a transposed auditory sequence has the peculiar characteristic of being composed of completely different sounds compared to the original sequence but keeping the same intervals between tones. The perception is therefore of a very similar sequence which is transposed to higher or lower pitches, as confirmed by the dissimilarity ratings reported in our study. Remarkably, the brain showed a very coherent behavior since its negative activity was exactly in between the conditions where the tones were exactly the same or completely different. This suggests that the recognition is a process where incoming temporal information is progressively matched with the previously stored memory trace. In this case, the transposed sequences were categorized at the same time as partially known and partially new sequences. Future studies may investigate whether the brain representation of same, transposed, and different melodies differs only for the intensity of the negative neural response or if different brain networks are involved.

Another key result emerged in our study is the difference between young and older adults in relation to the brain data. In fact, a large significant difference was observed when comparing the brain activity of these two categories of participants. Overall, the older adults showed a strongly reduced brain activity during the recognition and evaluation of the second sequence. This is even more interesting when considering that the magnitude of this difference was stronger than the one emerged when looking at the behavioral performance of the two age groups. This indicates the remarkable potential of neurophysiology of the auditory system to evaluate the neural efficiency and potentially health status of the neural system of older adults.

The reduction in terms of brain activity is coherent with several studies which reported a decreased ERP amplitude in older compared to young adults. For instance, Gajewski and Falkenstein ^27^ found that in the older compared to young adults the frontocentral N200 and the P300a and P300b were delayed and attenuated, especially when performing a two-back memory task. Similarly, Toledo, Manzano, Barela and Kohn ^64^ reported that during passive ankle movement, older adults had N100 characterized by lower amplitude and longer latency compared to young participants. Furthermore, in a suppression of visual information task ^7^, correct performances in young adults were associated with a robust modulation of alpha power. Conversely, this effect was not observed in older adults. Finally, in the linguistic domain, Federmeier, McLennan, De Ochoa and Kutas ^4^ demonstrated that older compared to young adults had a reduced neural efficiency when processing spoken language and recognizing familiar words. Similarly, previous investigations showed altered brain signatures associated with ageing when looking at resting state. For instance, Scally, Burke, Bunce and Delvenne ^65^ showed a reduction in the global EEG alpha power and functional connectivity in resting state for older adults. Likewise, Chow, Rabi, Paracha, Hasher, Anderson and Alain ^66^ reported reduced functional connectivity in alpha band during resting state when comparing older versus young adults. They highlighted especially the impaired connectivity between medial prefrontal cortex (mPFC) and several brain regions forming the default mode network. Moreover, they revealed that decreased functional connectivity between mPFC and temporal cortices was associated with reduced executive functions and semantic fluency in older adults. Along this line, another study ^67^ confirmed that older adults presented a reduced power for the alpha peak frequency and an increased power for beta, compared to young participants. Interestingly, the authors reported that better prossacade performance was associated with lower beta power in young participants only, suggesting that the increased beta power in older adults was a sign of cognitive decline.

In conclusion, our results showed the neural correlates of short-term recognition of auditory sequences, revealing a strong negativity associated with the recognition of previously heard sequences. In addition, we reported a very large reduction in the brain activity of older compared to young adults when doing the task, even when their behavioral performance was not altered. Taken together, our results reached a very strong statistical significance and effect size, suggesting that they are reliable and solid. However, since we used the EEG, we did not perform the source reconstruction of the recorded brain activity. Thus, we encourage future research to use MEG and MRI to compute the source reconstruction and provide additional information about the specific brain networks recruited for short-term recognition of auditory sequences. In addition, based on the several evidences which connected cognitive abilities with musical skills, musical perception, and brain structural and functional individual differences ^68-85^, future research should investigate the relationship between neural data underlying short-term memory for musical sequence and cognitive abilities such as fluid intelligence and WM. Finally, future studies may increase the sample size and provide further evidence on the impact of ageing on the brain activity. This would be of high importance for studying both successful ageing and pathological conditions such as dementia. In this light, we argue that the combination of auditory sequences, music, and fast-scale neurophysiology may represent a privileged solution to better understand the cognitive decline associated with ageing and potentially build novel tools to predict the probability of developing dementia.

## Acknowledgements

The Center for Music in the Brain (MIB) is funded by the Danish National Research Foundation (project number DNRF117).

LB is supported by the Carlsberg Foundation (CF20-0239), Lundbeck Foundation (Talent Prize 2022), Center for Music in the Brain, Linacre College of the University of Oxford (Halsall Fund), Nordic Mensa Fund.

MLK is supported by Center for Music in the Brain and Centre for Eudaimonia and Human Flourishing, which is funded by the Pettit and Carlsberg Foundations.

We thank Ivana Tomasi and Giada Pappalardo for their assistance in part of the EEG data collection.

## Author contributions

MC and LB conceived the hypotheses and designed the study. MC, LB, MLK and PV recruited the resources for the study. MC collected the data. MC performed pre-processing and statistical analysis, with help from LB. LB, MLK and PV provided essential help to interpret and frame the results within the neuroscientific literature. MC and LB wrote the first draft of the manuscript. MC prepared the figures. All the authors contributed to and approved the final version of the manuscript.

## Data availability

The codes and the neurophysiological data related to the experiment will be made available upon reasonable request.

